# A Regulatory Axis for Tonotopic MYO7A Expression in Cochlear Hair Cells

**DOI:** 10.1101/2025.04.04.647241

**Authors:** Sihan Li, Sujin Jun, Ye-Ri Kim, Ayathi Gogineni, Franklin Lee, Chul Hoon Kim, Un-Kyung Kim, Anthony W. Peng, Jung-Bum Shin

## Abstract

*Myo7a*, a gene mutated in Usher syndrome and non-syndromic deafness, encodes an unconventional myosin essential for hair cell function. Our previous work revealed that cochlear hair cells express distinct *Myo7a* isoforms with unique spatial and cell type-specific patterns. The canonical isoform (*Myo7a-C*) and a novel isoform (*Myo7a-N*) are co-expressed in outer hair cells (OHCs) but exhibit opposing tonotopic gradients, while inner hair cells (IHCs) primarily express *Myo7a-C*. These isoforms arise from distinct transcriptional start sites, indicating separate regulatory inputs. Here, we identify an intronic cis-regulatory element, *EnhancerA*, essential for tonotopically graded *Myo7a* expression. *EnhancerA* deletion reduces MYO7A protein levels in a tonotopically varied manner, disrupts hair bundle morphogenesis, alters OHC mechanotransduction, and leads to hair cell degeneration and hearing loss. We further identify SIX2, a tonotopically expressed transcription factor that may interact with *EnhancerA* to regulate *Myo7a-N* in OHCs. These findings define a cis-trans regulatory axis critical for isoform-specific *Myo7a* expression and cochlear function.

**Significance:** Cochlear hair cells rely on the molecular motor MYO7A for mechanosensory function. Our previous study revealed that Myo7a isoforms are differentially expressed in auditory hair cells, however, the mechanisms regulating this isoform-specific expression remain unclear. In this study, we identify a cis-regulatory element, *EnhancerA*, that governs MYO7A expression in a tonotopically graded manner. Furthermore, we demonstrate that the transcription factor SIX2 plays a role in regulation of MYO7A, and is essential for hair cell maintenance. These findings establish a mechanistic link between cochlear position and protein isoform diversity, highlighting how hair cells adapt to frequency-specific mechanical demands. Importantly, the tonotopic regulatory function of *EnhancerA* also offers new avenues for targeted gene therapy in auditory disorders.

## 3. Introduction

Sensory hair cells of the inner ear rely on the unconventional myosin motor protein MYO7A for proper development^1–4^, maintenance of hair bundles^5,6^, and mechanotransduction^7^. Loss-of-function mutations in the *Myo7a* gene cause both syndromic and non-syndromic forms of hereditary hearing loss. Notably, mutations in *MYO7A* underlie Usher syndrome type 1B, a disorder characterized by congenital deafness, vestibular dysfunction, and progressive vision loss^1,2,8^. In non-syndromic cases, *Myo7a* mutations account for a significant proportion of autosomal recessive deafness^9,10^. Given its central role in auditory function, MYO7A has emerged as a key target for inner ear gene therapy.

Previous studies have identified several *cis*-regulatory elements near the *Myo7a* locus^11,12^, including a proximal promoter located upstream of the canonical transcriptional start site^7,11^. This promoter has been incorporated into gene therapy vectors and can drive robust expression in inner hair cells (IHCs). However, its activity in outer hair cells (OHCs) is weak or inconsistent, limiting its effectiveness for pan-hair cell targeting^13,14^. Regulatory sequences derived from non-mammalian species, such as conserved elements in zebrafish^12^, have demonstrated hair cell-specific activity in transgenic models, but their correspondence to mammalian regulatory mechanisms and isoforms remains poorly understood.

Our previous work showed that cochlear hair cells express multiple *Myo7a* isoforms, including the genomically annotated canonical isoform (*Myo7a-C*), the short isoform (*Myo7a-S*), and more recently, a novel isoform (Myo7a-N) using long-read RNA sequencing from cochlear cDNA^15^. Hair cells lacking both *Myo7a-C* and *Myo7a-N* exhibit near complete loss of MYO7A levels at P5 and severe hair bundle disorganization with profound hearing loss by P20^15^, indicating that *Myo7a-S* is minimally expressed and that *Myo7a-C* and *Myo7a-N* are the predominant MYO7A isoforms in cochlear hair cells. Notably, *Myo7a-C* and *Myo7a-N* exhibit distinct cell type-specific and tonotopic distributions. *Myo7a-C* and *Myo7a-N* are both expressed in OHCs, but exhibit opposing gradients along the cochlear tonotopic axis: *Myo7a-N* expression increases toward the cochlear base, while *Myo7a-C* is enriched toward the apex. In contrast, *Myo7a-C* is predominant in IHCs across all tonotopic regions ^10,15^.

*Myo7a* isoforms arise from separate transcriptional start sites^16,17^, suggesting differential promoter usage and distinct trans-regulatory controls. Here, we identify a previously uncharacterized intronic enhancer, *EnhancerA*, that is essential for tonotopically graded expression of *Myo7a* isoforms in both IHCs and OHCs. Deletion of *EnhancerA* leads to a progressive loss of MYO7A protein toward the cochlear base, impaired hair bundle morphogenesis, altered mechanotransduction, hair cell degeneration, and hearing loss. We further identify SIX2, a transcription factor with tonotopically graded expression, as a potential trans-regulator that may interact with *EnhancerA* to selectively control *Myo7a-N* expression in OHCs. Loss of SIX2 does not produce immediate or severe phenotypes, but in aging mice, reduces *Myo7a-N* expression, leads to disorganization of OHCs, and causes progressive auditory dysfunction. Together, these findings define a previously unrecognized *cis-trans* regulatory axis that governs spatial and isoform-specific expression of *Myo7a* in the cochlea and highlight new molecular mechanisms essential for hair cell function and therapeutic intervention.

## 3. Results

### 3.1. *EnhancerA* is required for tonotopically graded expression of *Myo7a*

We previously showed that *Myo7a* is expressed in multiple isoforms within the cochlea, each exhibiting distinct tonotopic expression patterns^16,17^. Importantly, these isoforms are generated with different transcriptional start sites (**Fig. 1A**), suggesting that their regulation may involve distinct combinations of *cis*- and *trans*-acting elements. Previous studies have identified multiple enhancers and promoters of *Myo7a*, yet none have been shown to support broad and robust expression across all cochlear hair cells^13,14^. We therefore hypothesized that additional regulatory elements may contribute to the spatially patterned and cell type-specific expression of *Myo7a*.

**Figure 1:**
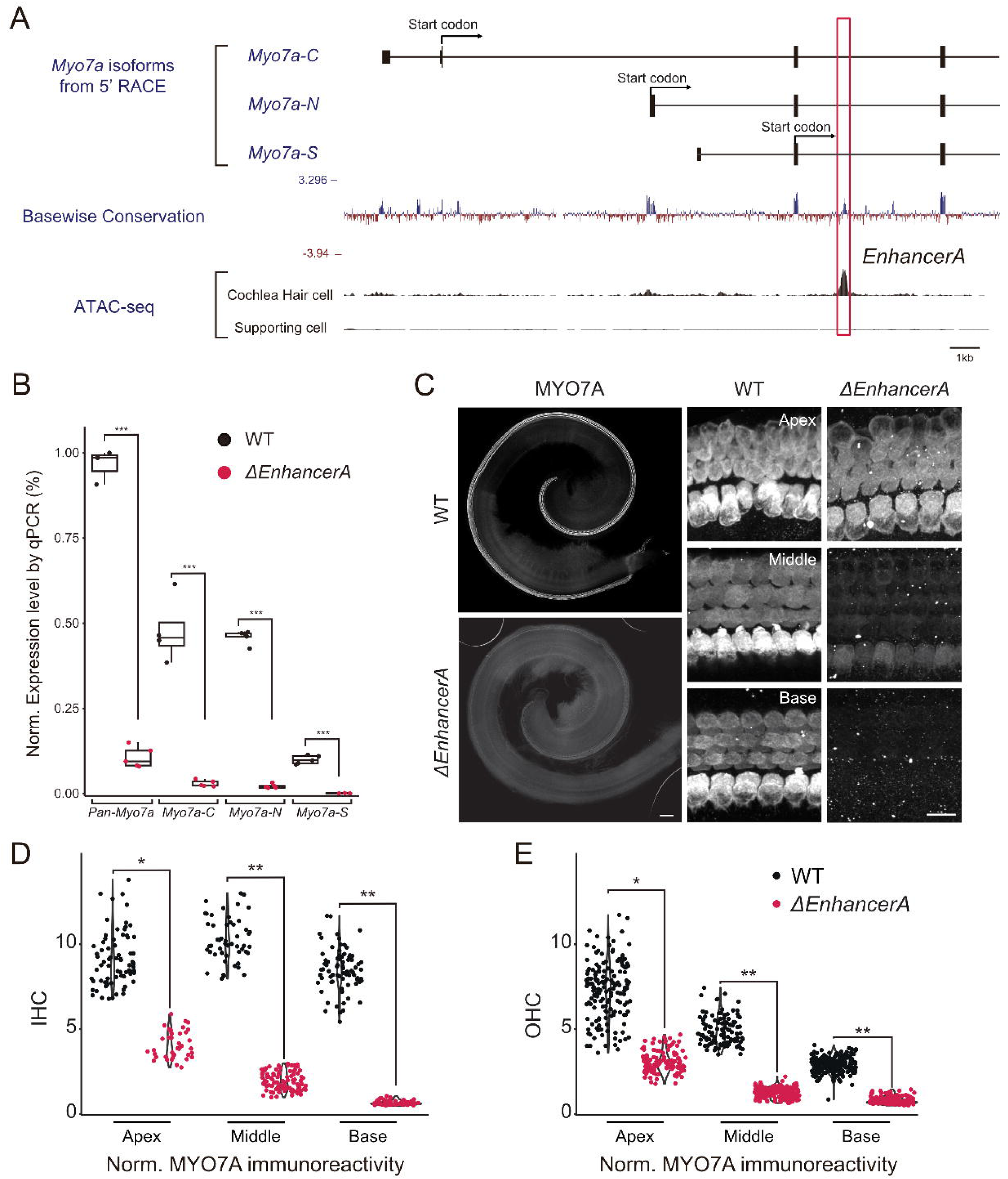
EnhancerA is required for tonotopically graded expression of Myo7a. **A**. Schematic indicating the transcriptional start sites of *Myo7a* isoforms. ATAC-seq tracks are derived from a publicly available dataset. *EnhancerA* is an around 400 bp open chromatin region that locates in the third intron of *Myo7a-N*, which is indicated with red bracket (GRCm38/mm10: chr7: 98105422-98105821). This region is specifically knocked out in Δ*EnhancerA* mice without affecting exons. Three major isoforms of Myo7a with distinct transcriptional start sites are demonstrated in the “*Myo7a*-isoform by 5’ RACE”. Basewise conservation shows multiple alignments of 35 vertebrate species as measurements of evolutionary conservation. **B**. Normalized expression level of *Myo7a* isoforms measured by qPCR on P0 cochlear cDNA. *Myo7a* expression level is normalized to *Gapdh* and then log-transformed. (WT vs. Δ*EnhancerA* mice t-test p-values of different isoforms: *pan-Myo7a*: 1.30e-4, *Myo7a-C*=2.60e-3, *Myo7a-N*=1.32e-07, *Myo7a-S*=5.59e-05. Number of Cochlea: WT=8, Δ*EnhancerA*=8. Pipetting repetition=5 per group). Boxplots show medians, 25th, and 75th percentiles as box limits and minima and maxima as whiskers. **C**. Immunofluorescence images of whole mount Δ*EnhancerA* cochlea at P7. Compared to WT cochlea, Δ*EnhancerA* mice showed tonotopic reduction of MYO7A immunoreactivity, with a stronger reduction in the basal region and less reduction at the apical region. (Scale bar: left: 100 µm, right 10 µm). **D, E.** Quantification of MYO7A immunoreactivity in IHCs and OHC at P7. MYO7A signals were normalized to MYO6. (Normalization method: MYO7A signal in the cell body is divided by F-actin signal in the cell body. T-test on the average normalized MYO7A immunofluorescence signal of each cochlea. Number of cochlea: WT: 4; Δ*EnhancerA*: 4. T-test p-value between WT and Δ*EnhancerA*: IHC: base: 5.63e-3; middle: 1.10e-3; apex: 1.43e-2. OHC: base: 1.77e-3; middle: 8.17e-3; apex: 2.89e-2).

To identify such elements, we reanalyzed publicly available ATAC-seq datasets from purified cochlear hair cells and supporting cells^18^. This analysis revealed a prominent, hair cell-specific peak of chromatin accessibility located within the third intron of the *Myo7a* gene, approximately 400 bp in length (**Fig. 1A**, genomic location provided in the figure legend). This peak represents the most accessible chromatin region within a 1Mb window surrounding *Myo7a*, consistent with a potential regulatory function. We designated this element *EnhancerA*.

To assess its functional role *in vivo*, we generated a mouse line harboring a 399 bp deletion targeting *EnhancerA* (Δ*EnhancerA* mice). Quantitative PCR analysis of cochlear cDNA from postnatal day 0 (P0) Δ*EnhancerA* mice revealed significantly reduced expression of all known *Myo7a* isoforms relative to wild-type (WT) controls, including *Myo7a-C* and *Myo7a-N*, and *Myo7a-S*. This result indicates that *EnhancerA* is essential for robust *Myo7a* transcription but not selective to *Myo7a* isoforms (**Fig. 1B**).

Strikingly, immunofluorescence analysis of MYO7A protein levels in Δ*EnhancerA* mice at P7 revealed a pronounced and spatially patterned reduction in both IHCs and OHCs (**Fig. 1C**, quantified for IHCs and OHCs in **Fig. 1D, E**). In the basal cochlea, MYO7A protein was nearly undetectable, indicating a severe transcriptional or post-transcriptional deficit. In contrast, the apical cochlear region retained robust MYO7A expression, though levels were reduced by approximately 50% compared to WT controls. Notably, reduced MYO7A levels are observed in both OHCs and IHCs, consistent with our qPCR results showing that all MYO7A isoforms are affected by *EnhancerA* deletion. MYO7A expression is broadly comparable across tonotopic regions in WT controls. The progressive base-to-apex gradient of MYO7A loss observed upon *EnhancerA* deletion suggests that *EnhancerA* functions as a key cis-regulatory element that supports robust, spatially biased MYO7A expression along the cochlear axis. Importantly, this regulatory effect appears to act without isoform selectivity.

### 3.2. *EnhancerA* deletion impairs hair bundle development

To evaluate the developmental consequences of *EnhancerA* deletion, we performed scanning electron microscopy (SEM) on Δ*EnhancerA* cochleae at P7, a stage when hair bundle maturation is largely complete. In the most basal cochlear region (“hook” region), where MYO7A loss was the most severe, we observed marked disorganization of hair bundles in both IHCs and OHCs (**Fig. 2A, B**). Defects included the near-complete absence of the shortest row of stereocilia, a reduced number of middle and tallest row stereocilia, and disorganized stereocilia clusters. The clustering phenotype suggests impaired formation or maintenance of interstereociliary links, a process dependent on MYO7A function^19–21^.

**Figure 2:**
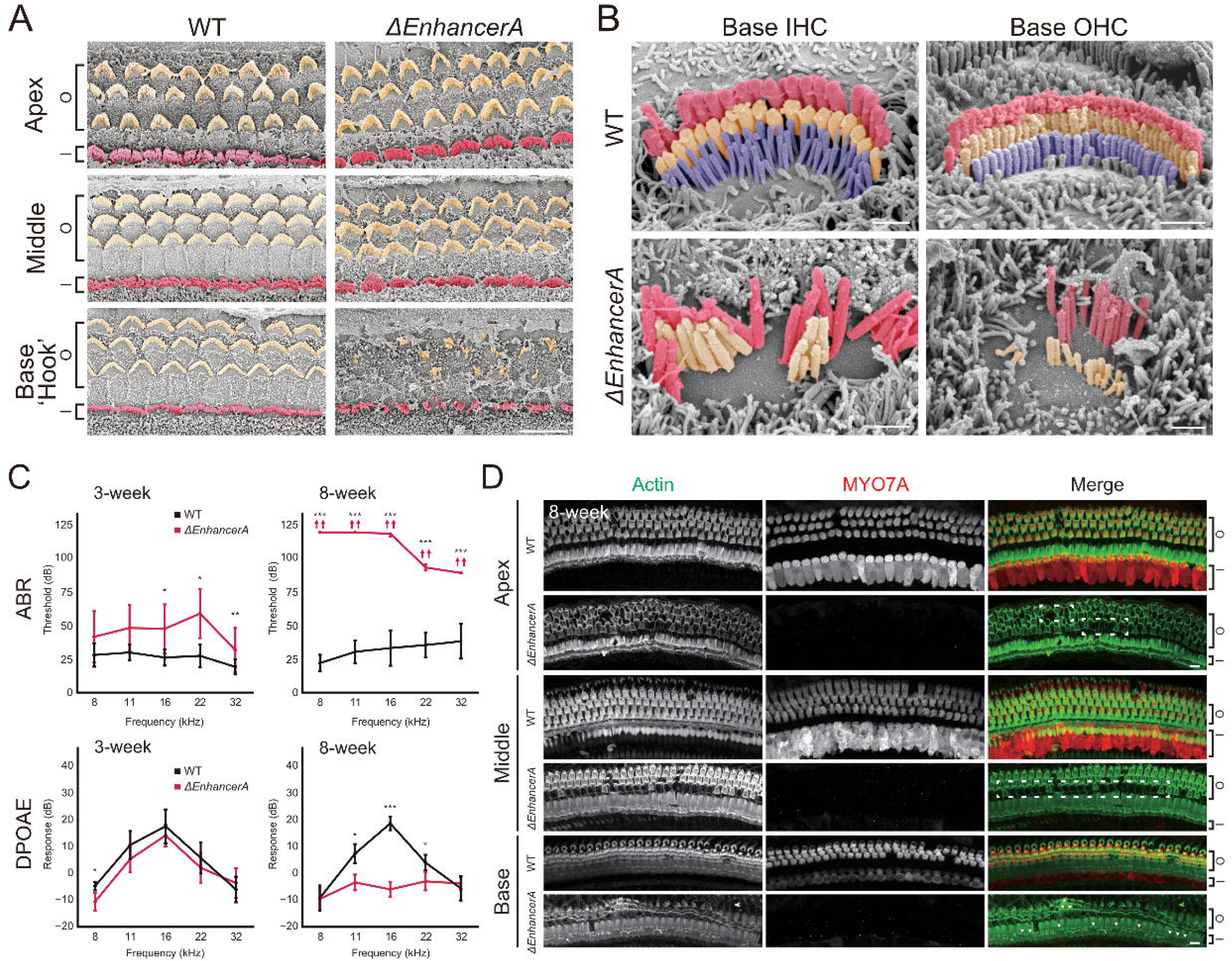
Hair cell degeneration and hearing loss of Δ*EnhancerA* mice. **A**. Scanning electron microscopy (SEM) images of WT and Δ*EnhancerA* cochleae at P7. Δ*EnhancerA* hair cells display disorganized hair bundles near the base of the cochlea. (Scale bar: 10 µm). **B**. Higher-magnification SEM images of WT and Δ*EnhancerA* cochleae at P7 reveal detailed structural abnormalities in Δ*EnhancerA* hair bundles. (Scale bar: 1 µm). **C**. ABR thresholds and DPOAE response of WT and Δ*EnhancerA* at 3 weeks and 8 weeks of age. Δ*EnhancerA* mice exhibit mild hearing loss at 3 weeks, which progresses to profound hearing loss by 8 weeks. (ABR ANOVA p-value: 3-week=1.25e-08, 8-week<2e-16. Number of animals: WT: 3-week=8, 8-week=9; Δ*EnhancerA*: 3-week=8, 8-week=5. DPOAE p-value: 3-week=0.0060, 8-week=1.12e-11. Number of animals: WT: 3-week=8, 8-week=10; Δ*EnhancerA*: 3-week=7, 8-week=5). Line-plots show means of the ABR or DPOAE response. Standard error is presented as error bars. **D**. Immunofluorescence images of WT and Δ*EnhancerA* mice at 8 weeks. MYO7A expression is nearly absent across all cochlear regions in Δ*EnhancerA* mice, accompanied by extensive hair cell loss. (Scale bar: 10 µm. Asterisk labels the remaining OHCs at basal regions of Δ*EnhancerA* mice. Arrowheads label the IHCs loss at basal area. White dashed box highlights the OHC loss in middle and apical turn of cochlea).

In contrast, hair cells in more apical regions, where residual MYO7A protein remained, displayed largely normal bundle morphology resembling that of WT controls (**Fig. 2A**). These findings indicate that *EnhancerA* is required for proper stereocilia development in a spatially graded manner: based on the observation of impaired hair bundles in the Δ*EnhancerA* “hook” region, it is likely that the reduced level of MYO7A in this region is insufficient to support proper hair bundle development or organization. In contrast, in the middle and apical regions, MYO7A levels are reduced to a lesser extent, allowing hair bundles to maintain a WT-like morphology.

### 3.3. Hair cell loss and hearing loss in Δ*EnhancerA* mice

To determine whether these structural defects lead to auditory dysfunction, we measured auditory brainstem responses (ABRs) and distortion product otoacoustic emissions (DPOAEs). At 3 weeks of age, Δ*EnhancerA* mice exhibited mild to moderate hearing loss, with significant threshold shifts at mid-to-high frequencies (**Fig. 2C**), consistent with the basal-to-apical gradient of MYO7A depletion. Lower frequencies also showed a statistically insignificant trend toward elevated ABR thresholds, likely reflecting that the residual MYO7A levels are insufficient to fully support optimal hair cell function. Distortion product otoacoustic emissions (DPOAEs) were largely unaffected at this stage. However, by 8 weeks, Δ*EnhancerA* mice exhibited profound hearing loss across all frequencies, accompanied by the near-complete loss of DPOAEs (**Fig. 2C**).

To further explore the cellular basis of this rapid progression, we performed immunofluorescence analysis of MYO7A and hair cell morphology at 8 weeks. By this stage, MYO7A expression was nearly absent throughout the cochlea in Δ*EnhancerA* mice (**Fig. 2D**), and was accompanied by extensive degeneration of hair cells in the basal turn, with partial loss observed in the middle and apical turn (**Fig. 2D**). These results suggest that *EnhancerA* is not only essential for the early developmental expression of *Myo7a*, but also becomes increasingly critical for maintaining *Myo7a* expression and hair cell survival as the cochlea matures.

Together, these findings demonstrate that *EnhancerA* plays a critical role in regulating MYO7A expression in a tonotopically graded manner. Its deletion disrupts hair bundle morphogenesis, compromises hair cell maintenance, and impairs long-term auditory function, initially affecting high-frequency regions and progressively extending toward lower-frequency areas with age. Notably, the highest ABR frequency tested was 32 kHz, which corresponds to the mid-basal region of the cochlea. Therefore, functional deficits in the more basal, higher-frequency region may be underestimated in our measurements.

### 3.4. Deletion of *EnhancerA* causes reduced resting open probability of MET channels in a tonotopically graded manner

Our previous work demonstrated that isoform-specific deletion of *Myo7a-C* results in approximately 80% reduction of MYO7A protein in IHCs, leading to decreased resting open probability and delayed activation kinetics of MET currents^17^. These findings established a role for MYO7A-C in maintaining resting tension on the MET complex in IHCs. However, the contribution of MYO7A to MET function in OHCs was not characterized in our previous study. The Δ*EnhancerA* mouse model presents an opportunity to address this gap. Unlike the isoform-specific KO, Δ*EnhancerA* mice exhibit reduced expression of all known *Myo7a* isoforms in both IHCs and OHCs. Furthermore, MYO7A loss occurs in a tonotopically graded manner, allowing us to correlate spatial differences in protein levels with MET channel properties, independent of isoform bias.

To assess MET function, we recorded transduction currents from OHCs in the basal and apical cochlea of Δ*EnhancerA* homozygous mice (*EA^−/−^*) at postnatal day 4-5 (P4-5), using fluid jet stimulation and high-speed imaging to monitor bundle deflections. Considering that the “hook” region of Δ*EnhancerA* mice displays disorganized hair bundles, we first confirmed that stereocilia in the basal region preceding the “hook” region exhibit WT-like development, thereby avoiding potential confounds arising from disrupted hair bundle morphology (**Supplementary Fig. 1A**). All recordings were restricted to regions with intact stereocilia morphology from the mid-basal to apical sections of the cochlea.

Apical OHCs, where MYO7A expression is reduced by only 50%, exhibited no significant differences in resting open probability, peak current, or adaptation compared to heterozygous controls (*EA^+/+^*) (**Fig. 3A, B, E, F, Supplementary Fig. 2B**). However, in basal OHCs, where MYO7A levels are strongly reduced but bundles are still intact, sinusoidal stimulation revealed a significant decrease in the resting open probability of MET channels, while peak current amplitudes remained unchanged (**Fig. 3C, D**). Similarly, step-like stimuli confirmed a reduction in resting open probability, with no detectable changes in current amplitude or adaptation kinetics at P4-P5 (**Fig. 3E, G; Supplementary Fig. 2B**). Collectively, these results confirm our previous study that MYO7A levels are critical for maintaining resting tension on the MET apparatus^17^.

**Figure 3:**
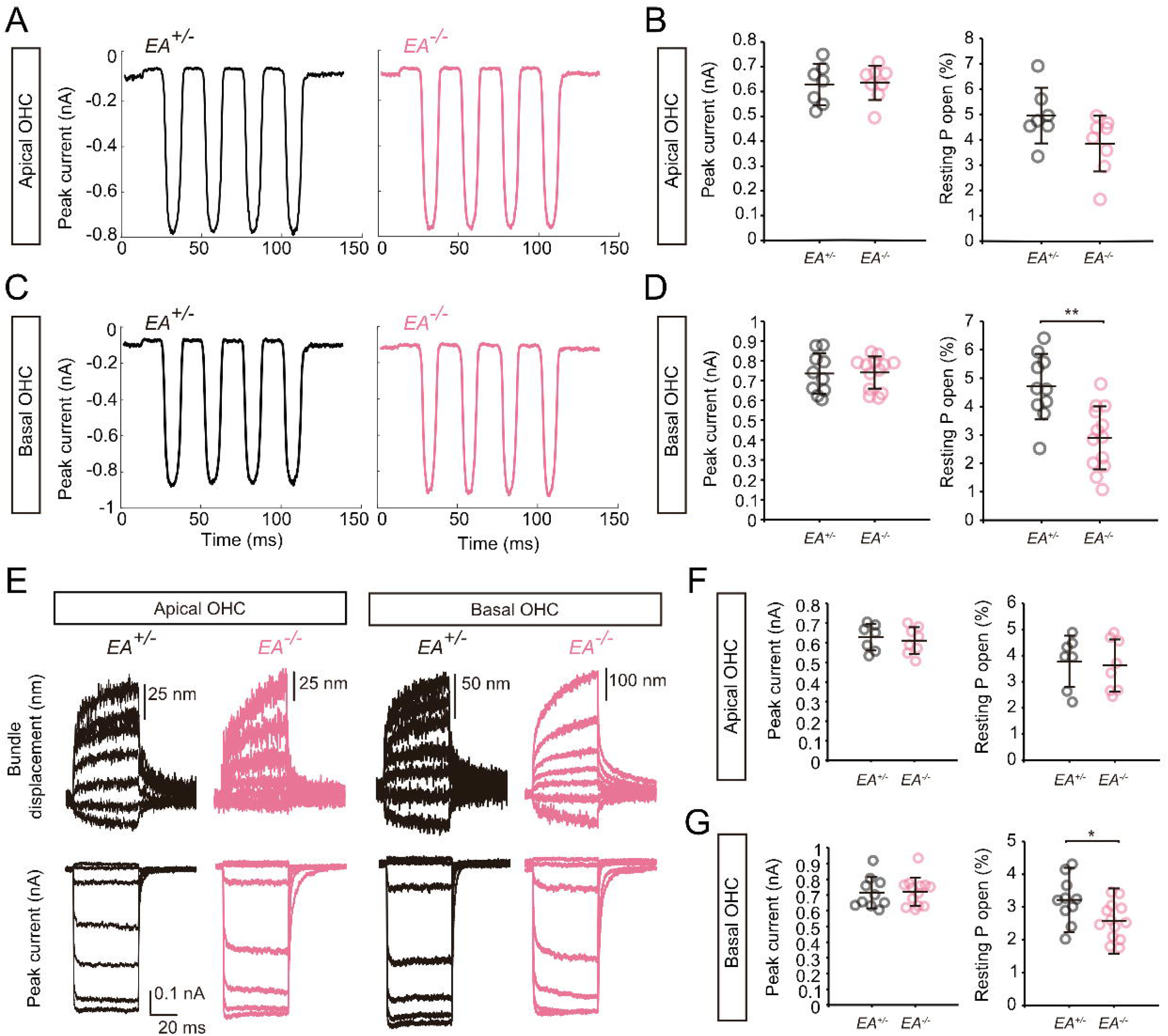
Reduction of resting open probability of Δ*EnhancerA* OHC. **A**. Evaluation of the MET current of apical OHCs from *EA^+/+^* (black) and *EA^−/−^* (pink) at P4-P5. MET currents are evoked by sinusoidal stimulation. **B**. Summary plots of the peak current and P_o_ observed from apical OHCs. Number of cells (animals): *EA^+/+^* 8 (8), *EA^−/−^* 7 (7). **C**. Evaluation of the MET current of basal OHCs from *EA^+/+^* and *EA^−/−^* at P4-P5. MET currents are evoked by sinusoidal stimulation. **D**. Summary plots of the peak current and P_o_ observed from basal OHCs. ** p ≤ 0.01. Number of cells (animals): *EA^+/+^* 13 (13), Basal *EA^−/−^* 10 (10). **E**. Comparison of MET currents in apical and basal turns of *EA^+/+^* and *EA^−/−^* mice measured using step-like force stimuli. Hair bundle displacement traces are shown at the top, and evoked peak MET currents are shown at the bottom. **F, G.** Summary plots of the peak current and resting open probability for apical (**F**) and basal (**G**) turn outer hair cells. * p ≤ 0.05. Number of cells (animals): Apical *EA^+/+^* = 8 (8), Basal *EA^+/+^* = 13 (13), Apical *EA^−/−^* = 7 (7), Basal *EA^−/−^* = 10(10). (For panel B, D, F and G, whiskers of the plot represent mean ± SD).

### 3.5. SIX2 is tonotopically expressed in the cochlea

While *EnhancerA* confers a tonotopically graded expression pattern on *Myo7a* expression, its deletion disrupts both *Myo7a-C* and *Myo7a-N* isoforms, indicating that additional mechanisms are involved in mediating isoform-specific regulation. To identify potential trans-regulatory factors, we queried the Unbind2021^22^ and ReMap2022^23^ ChIP-seq databases and discovered a prominent SIX2 binding site centrally located within *EnhancerA*^24,25^ (**Fig. 4A, B**). SIX2 is a homeodomain transcription factor known for its roles in kidney^26–29^, craniofacial skeleton development^30^. Although haploinsufficiency of SIX2 has been associated with mild high-frequency hearing loss in humans^31^, its function in auditory and vestibular hair cells remains largely uncharacterized.

**Figure 4:**
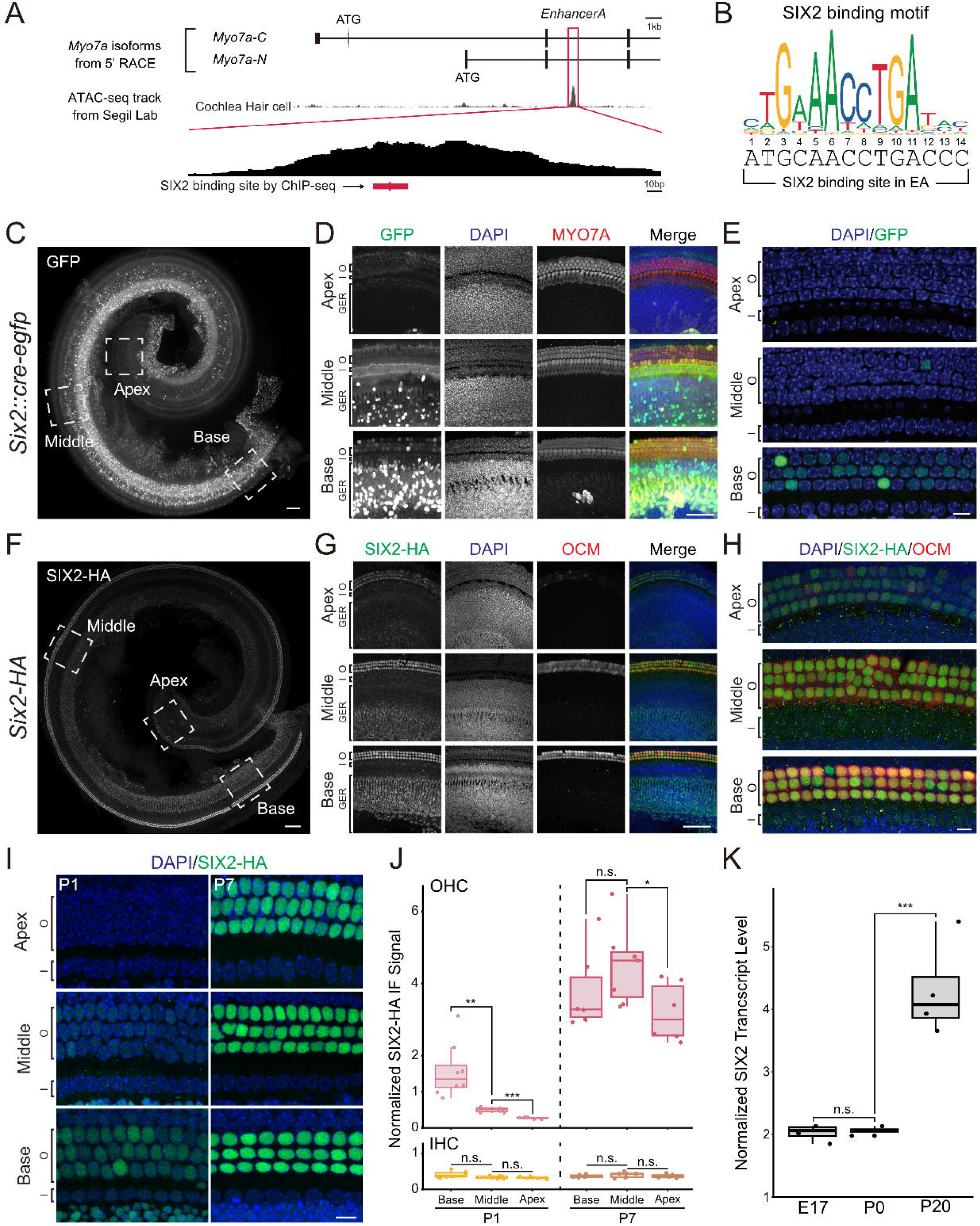
Tonotopic expression of SIX2 in cochlea. **A**. The diagram demonstrating the SIX2 binding site in the *EnhancerA* tested by published ChIP-seq data. **B**. Human SIX2 binding motif reported by JASPAR database. The DNA sequence of SIX2 binding site in the *EnhancerA* is provided below the binding motif. **C**. Whole mount immunofluorescence image of *Six2::cre-egfp* cochlea at P5. GFP is tonotopically expressed in the cochlea, with a higher expression level at base and decrease toward apex. (Scale bar: 100 µm). **D**. Immunofluorescence image of *Six2::cre-egfp* cochlea at base, middle and apex region. GFP was observed in the OHCs and GER cells. (Scale bar: 10 µm. “I”: IHCs; “O”: OHCs; “GER”: greater epithelial ridge cells). **E**. immunofluorescence image of *Six2::cre-egfp* hair cell regions at base, middle and apex region. (Scale bar: 10 µm). **F**. Whole mount immunofluorescence image of *Six2-HA* cochlea mice at P7. Similar with *Six2::cre-egfp* cochlea, SIX2 is also observed to be expressed tonotopically. (Scale bar: 100 µm). **G**. immunofluorescence image of *Six2-HA* cochlea at base, middle and apex region. HA-tag signals were observed in the OHCs and GER cells. Oncomodulin (OCM) is stained as OHC marker (Scale bar: 10 µm). **H**. immunofluorescence image of *Six2-HA* hair cell regions at base, middle and apex region. (Scale bar: 10 µm). **I**. Representative SIX2-HA immunostaining images at P0 and P7. Scale bar = 10 µm. **J**. Quantification of SIX2-HA fluorescence signal at P0 and P7. At P0, SIX2 is tonotopically expressed in OHCs, with the highest levels in the basal turn of the cochlea. SIX2 expression is significantly increased across all cochlear regions at P7. In contrast, SIX2 expression in IHCs remains low at both P0 and P7. (Normalization method: SIX2-HA signals are collected in the cell nuclei and then normalized to DAPI. Number of cochleae: P0 = 8; P7 = 6. T-test p-values: P0: IHC: base vs. middle = 0.060; middle vs. apex = 0.62; OHC: base vs. middle = 5.00e-3; middle vs. apex = 5.58e-6. P7: IHC: base vs. middle = 0.36; middle vs. apex = 0.45; OHC: base vs. middle = 0.28; middle vs. apex = 0.033. **K**. Transcript levels of Six2 measured at E17, P0, and P20. (Normalization method: Cq value of SIX2 is first normalized to Cq value of Gapdh and then log-transformed. p-values: E17 vs. P0 = 0.66; P0 vs. P20 = 9.75e-3).

First, we investigated the expression pattern of SIX2 in the mouse cochlea. Previously, it has been reported that a widely used commercially available SIX2 antibody also interacts with SIX1^24^. To circumvent this, we utilized the *Six2::cre-egfp* mouse line and visualized EGFP expression as a proxy for SIX2 activity. At P5, we observed strong SIX2 driven EGFP signal in OHCs and a subset of supporting cells within the greater epithelial ridge (GER), with comparatively weaker expression in IHCs (**Fig. 4C–E**). Notably, *Six2::cre* activity displayed a tonotopic gradient, highest in the basal cochlear turn and tapering toward the apex, mirroring the distribution of *Myo7a-N* isoform expression. This parallel pattern suggests a potential regulatory role for SIX2 in modulating *Myo7a-N* expression in the cochlea.

To confirm endogenous SIX2 expression and define its subcellular localization, we generated a *Six2-HA* knock-in (KI) mouse line, tagging the C-terminus of the endogenous gene with an HA epitope. The HA tag did not grossly affect SIX2 function, since *Six2-HA* homozygous mice developed normally, while SIX2 deletion is known to cause perinatal lethality^26^. Immunostaining of these cochleae revealed a similar tonotopic pattern of SIX2-HA signal as observed in *Six2::cre-egfp* mice, with strong nuclear expression in OHCs and GER cells in the basal region, decreasing toward the apex (**Fig. 4F–H**). Consistent with *Six2::cre* activity, SIX2-HA expression was low in IHCs, supporting the idea that SIX2 acts as an OHC-specific transcription factor within the hair cell population. The nuclear enrichment of the HA signal further suggests that SIX2 is transcriptionally active in these cells.

To further characterize the expression pattern of SIX2 at different developmental stages in the cochlea, we extended our SIX2-HA immunostaining analyses to P1 and P7. At P1, SIX2 expression was detected in OHCs of the basal and middle turns, while minimal signal was observed in the apical turn (**Fig. 4I, J**). By P7, SIX2 expression in OHCs increased across all cochlear regions and became relatively uniform (**Fig. 4I, J**), indicating a loss of the initial tonotopic gradient. In contrast, SIX2 expression remained low in IHCs at both P1 and P7. These results suggest that SIX2 expression initiates at the basal end of the cochlea around P1 and subsequently expands to all cochlear regions by P7.

To determine whether SIX2 transcripts are expressed at embryonic and adult stages, we performed qPCR to assess SIX2 transcript levels at E17, P0, and P20 (**Fig. 4K**). SIX2 transcript levels showed low expression between E17 and P0. In contrast, SIX2 expression was significantly increased at P20, when hair cells are fully matured, suggesting a role for SIX2 in postnatal and adult hair cells.

Taken together, these findings identify SIX2 as a transcription factor that is preferentially expressed in OHCs relative to IHCs and whose expression is developmentally regulated, initiating around P0 with a transient tonotopic pattern that becomes uniform across the cochlea by P7.

### 3.6. Loss of SIX2 does not disrupt hair cell development but impairs hair cell maintenance

Based on the expression pattern of SIX2 in the cochlea, particularly its enrichment in OHCs, we hypothesized that SIX2 may play a critical role in OHC development or maintenance. We first tested whether SIX2 is required for OHC development. Because global deletion of SIX2 results in early postnatal lethality^26^, we generated a hair cell-specific SIX2 conditional knockout by crossing *Gfi1-cre* mice with *Six2^flox/flox^* mice (*Gfi1^cre/-^*; *Six2^flox/flox^*, hereafter referred to as, *Gfi1-Six2-cKO*). In this model, SIX2 is specifically deleted in hair cells during early embryonic stages, likely preceding the onset of endogenous SIX2 expression.

We first examined hair cell development at P7 using immunofluorescence microscopy. In the basal turn of the cochlea, no loss of IHCs or OHCs was observed in Gfi1-Six2-cKO mice, and OHCs maintained the normal three-row organization (**Fig. 5A, B**). Notably, MYO7A immunofluorescence signals in both basal IHCs and OHCs were comparable to those in control mice (**Fig. 5B**). Given that SIX2 is highly enriched in OHCs, we further examined hair bundle morphology in basal OHCs using SEM. This analysis revealed no disorganization of hair bundles, stereocilia degeneration, or other overt morphological abnormalities (**Fig. 5C**). Based on these results, we conclude that SIX2 is not required for early hair cell development, consistent with the postnatal-onset of its expression in OHCs.

**Figure 5:**
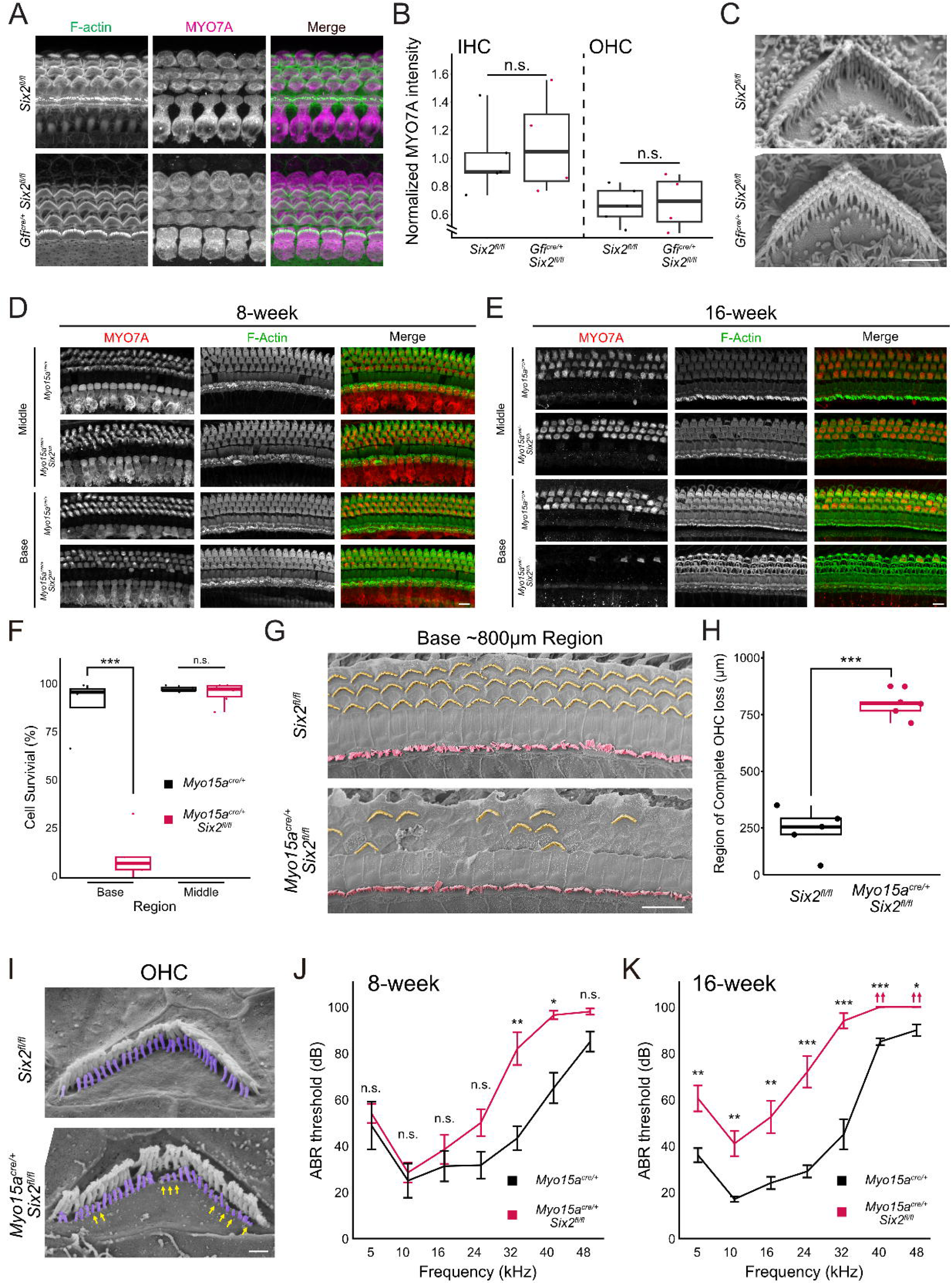
SIX2 is important for hair cell maintenance. **A**. Immunofluorescence images of *Six2^fl/fl^* mice and *Gfi^cre/+^; Six2^fl/fl^* mice at P7. (Scale bar: 10 µm). **B**. Quantification of MYO7A level in the cell body of basal turn. No significant MYO7A reduction was observed (Normalization method: MYO7A signal in the cell body is divided by F-actin signal in the cell body. Number of animals: *Six2^fl/fl^*: 4; *Gfi^cre/+^; Six2^fl/fl^*: 4. T-test p-value of *Six2^fl/fl^* vs. *Gfi^cre/+^; Six2^fl/fl^*: IHC: 0.66, OHC: 0.87). **C**. SEM images of *Six2^fl/fl^* and *Gfi^cre/+^; Six2^fl/fl^* OHCs at basal turn at P7. *Gfi^cre/+^; Six2^fl/fl^* mice do not show an obvious difference between control OHCs. (Scale bar: 1 µm). **D**. Representative image of 8-week control (*Myo15a^cre/+^*) and *Myo15a^cre/+^; Six2^fl/fl^* cochlea at middle and basal turns. (Scale bar: 10 µm). **E**. Representative image of 16-week control and *Myo15a^cre/+^; Six2^fl/fl^* cochlea at middle and basal turns. *Myo15a^cre/+^* induced conditional deletion of *Six2* results in OHC degeneration starting from basal turn. (Scale bar: 10 µm). **F**. Quantification of OHCs of control and *Myo15a^cre/+^; Six2^fl/fl^* cochlea at middle and basal turns at 16-week. (Number of animals: *Myo15a^cre/+^*: 3, *Myo15a^cre/+^; Six2^fl/fl^* = 3. T-test p-value: base = 1.85e-4; middle = 0.39.). **G.** SEM images of *Six2^fl/fl^* and *Myo15a^cre/+^*; *Six2^fl/fl^* cochleae at 10-week. Images were taken at approximately 800 µm from the basal end of the cochlea. (Scale bar: 10 µm). **H.** Quantification of the length of the cochlear region exhibiting complete OHC loss, measured from the beginning of basal end. (Number of animals: *Six2^fl/fl^*: 5; *Myo15a^cre/+^*; *Six2^fl/fl^*: 5. Average length: *Six2^fl/fl^*: 240.02 µm; *Myo15a^cre/+^*; *Six2^fl/fl^*: 791.80 µm. T-test p-value: 2.41e-05). **I.** SEM images of *Six2^fl/fl^* and *Myo15a^cre/+^*; *Six2^fl/fl^* OHCs in the basal turn at 10 weeks of age. The third row of stereocilia is highlighted in light blue. Degenerating stereocilia is marked by yellow arrows. (Scale bar: 1 µm). **J**. ABR test of *Myo15a^cre/+^*; *Six2^fl/fl^* mice at 8 weeks. Mild ABR threshold shift was observed in the high frequencies tested. (ANOVA p-value: 3.23e-06. Number of animals: *Six2^fl/fl^* = 4, *Myo15a^cre/+^*; *Six2^fl/fl^* = 10). **K**. ABR test of *Myo15a^cre/+^*; *Six2^fl/fl^* mice at 16 weeks. ABR threshold shift was observed in all of frequencies tested. (ANOVA p-value: 7.85e-15. Number of animals: *Six2^fl/fl^* = 6, *Myo15a^cre/+^*; *Six2^fl/fl^* = 10).

Based on the expression of SIX2 that starts around P0 and progressively increases into adult stages, we hypothesized that SIX2 contributes to hair cell maintenance. Since *Gfi1-cre* causes hearing loss in adult mice due to haploinsufficiency^32^, we chose *Myo15a-cre* to investigate the role of SIX2 in mature hair cells (*Myo15a^cre/+^*; *Six2^flox/flox^* mice hereafter referred to as *Myo15a-Six2-cKO*). Because *Myo15a-Cre* expression begins during the early postnatal period ^33^, a few days after the onset of SIX2 expression, this strategy enables selective deletion of SIX2 in hair cells after initial hair cell differentiation and development. At 8 weeks of age, *Myo15a-Six2-cKO* mice showed no apparent loss of hair cells (**Fig. 5D**). However, by 16 weeks, we observed progressive degeneration of OHCs, starting in the basal turn of the cochlea (**Fig. 5E-F**).

Because no OHC loss was observed in *Myo15a-Six2-cKO* mice at 8 weeks of age, whereas near-complete OHC loss was evident in the basal turn by 16 weeks, we sought to identify an intermediate stage to examine OHC hair bundle morphology at the basal turn and further assess the effects of SIX2 deletion on hair bundles. We first confirmed that 10-week, *Myo15a-Six2-cKO* mice exhibit increased OHC loss by quantifying the length of the organ of Corti with complete OHC loss, which was significantly longer compared with controls (∼800 µm in cKO vs. ∼250 µm in control) (**Fig. 5G-H**). In addition, surviving OHCs located approximately 800 µm from the basal end displayed degeneration of the third row of stereocilia (**Fig. 5I**).

OHC degeneration was accompanied by progressive hearing loss, with threshold shifts first emerging at mid-to-high frequencies by 8 weeks and expanding to all frequencies by 16 weeks (**Fig. 6J–K**). These findings suggest that SIX2 is required for the maintenance of hair cell integrity post-development, and its deletion leads to hair cell loss, hair bundle degeneration and hearing impairment.

**Figure 6:**
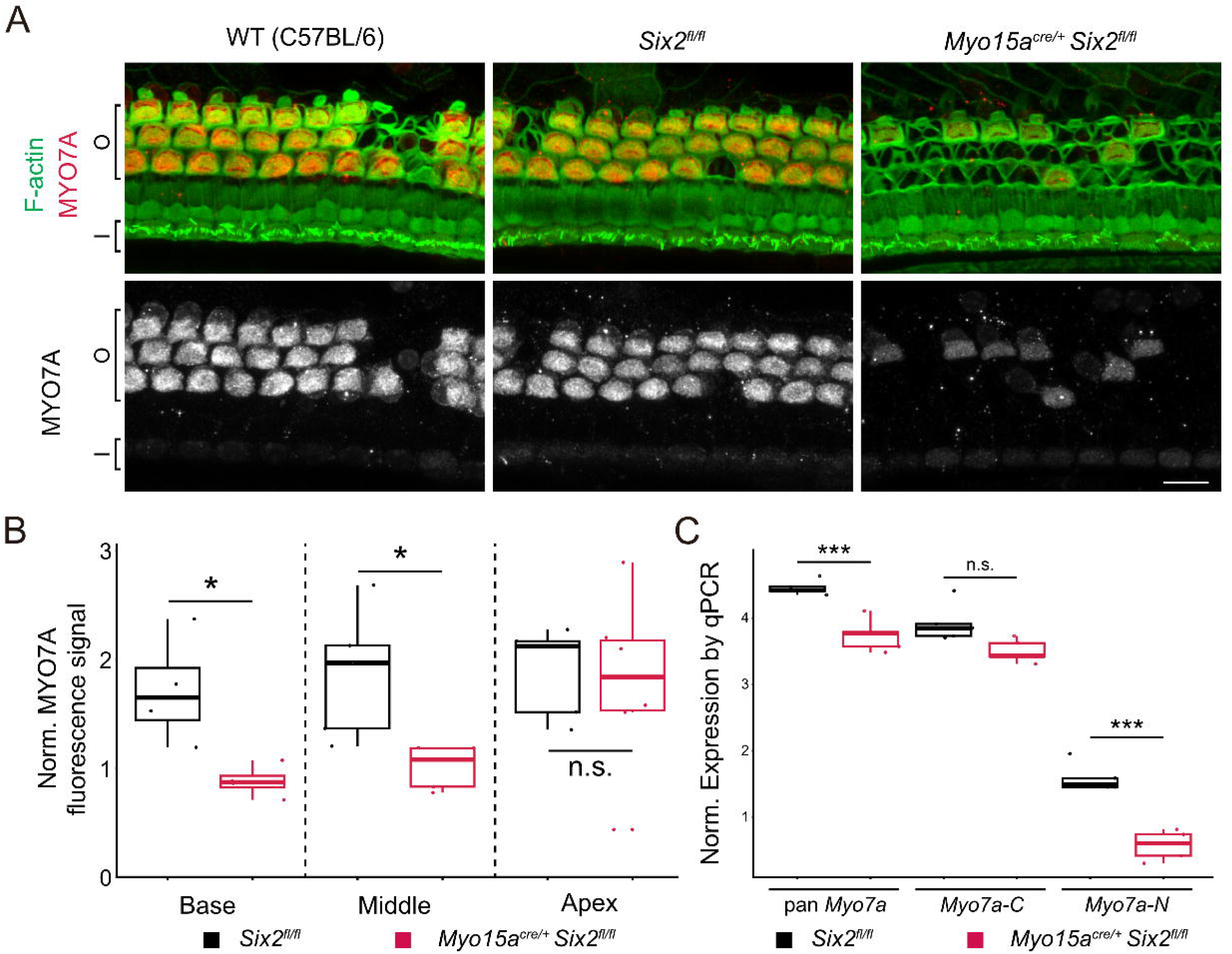
SIX2 regulates MYO7A expression. **A**. Representative OHC image of 10-week WT (C57BL/6), *Myo15a^cre/+^*; *Six2^fl/fl^* and *Six2^fl/fl^*cochlea at middle and basal turns. (Scale bar: 10 µm). **B**. Quantification of MYO7A level in the cell body of OHCs at 10-week. MYO7A is significantly reduced in basal and middle turn of the cochlea (Normalization method: MYO7A signal in the cell body is divided by F-actin signal in the cell body. Number of animals: *Six2^fl/fl^*: 5; *Myo15a^cre/+^*; *Six2^fl/fl^*: 5. T-test p-value of *Six2^fl/fl^* vs. *Myo15a^cre/+^*; *Six2^fl/fl^*: base: 0.039, middle: 0.030, apex: 0.80). **C**. Normalized expression level of *Myo7a* isoforms of *Myo15a^cre/+^*; *Six2^fl/fl^* and control *Myo15a^cre/+^* cochlear cDNA at P20. *Myo7a* expression level is normalized to *Gapdh* and then log-transformed. (T-test p-value: *pan-Myo7a* = 1.95e-4; *Myo7a-C* = 7.04e-2; *Myo7a-N* = 2.66e-3. Technical replication: 5 per group).

### 3.7. SIX2 regulates expression of *Myo7a*

Lastly, given that SIX2 is highly enriched in OHCs and exhibits a tonotopic expression pattern during early postnatal stages that correlates with *Myo7a-N*, we further tested the hypothesis that SIX2 regulates the expression of MYO7A protein and *Myo7a* transcripts. At 10 weeks of age, we observed a significant reduction of MYO7A levels in surviving OHCs in the basal turn of *Myo15a-Six2-cKO* mice. In addition to the basal turn, OHCs in the middle turn also showed a significant decrease in MYO7A signal, whereas the apical turn was not affected (**Fig. 6A, B**). Notably, we also observed an unexpectedly low expression level of MYO7A in IHCs. However, this low expression was also found in regular WT controls, indicating that the decreased MYO7A signal in IHCs is not caused by the *Six2^fl/fl^* genotype but instead reflects a normal effect of aging (**Fig. 6A**). Overall, this tonotopically restricted reduction of MYO7A correlates with the high-frequency hearing loss observed in *Myo15a-Six2-cKO* mice, suggesting that SIX2 regulates MYO7A expression preferentially in the high-frequency regions of the cochlea at 10 weeks of age.

We next analyzed *Myo7a* isoform expression using qPCR. We chose to perform this at postnatal day 20 (P20), reasoning that transcript changes will precede protein loss, and any transcript changes at this stage likely reflect regulatory effects rather than cell loss. Notably, expression of total *Myo7a* and the *Myo7a-N* isoform was significantly reduced in SIX2-deficient mice, whereas the *Myo7a-C* isoform was not significantly affected (**Fig. 6C**). These results suggest that SIX2 regulates *Myo7a* expression and preferentially influences the expression of MYO7A-N.

Together, these data reveal that SIX2 plays a crucial role in maintaining gene expression in mature hair cells, including the regulation of specific *Myo7a* isoforms, and that its loss compromises hair cell survival and auditory function.

## 4. Discussion

### *EnhancerA* orchestrates tonotopic *Myo7a* expression in cochlear hair cells

Cochlear hair cells express two *Myo7a* isoforms, *Myo7a-C* and *Myo7a-N*, with complementary tonotopic and cell type-specific expression^16,17^. While IHCs predominantly express *Myo7a-C*, OHCs co-express both isoforms in opposing spatial gradients. Furthermore, these two isoforms exhibit distinct ATPase activity, suggesting their roles in regulating MET properties. Thus, understanding how this expression diversity is regulated is expected to be important for decoding their distinct functional roles.

Here, we identify *EnhancerA*, a conserved intronic *cis*-regulatory element, as essential for maintaining robust *Myo7a* expression in both IHCs and OHCs. Deletion of *EnhancerA* during early postnatal development causes a progressive, tonotopically graded reduction in MYO7A transcript and protein level, culminating in near-complete loss by 8 weeks. This suggests that *EnhancerA* activity becomes increasingly critical as chromatin landscapes mature.

Importantly, *EnhancerA* deletion impacts both isoforms, indicating that it serves as a shared enhancer of isoform-selective promoters. The widely used canonical promoter upstream of *Myo7a-C* is highly active in IHCs but weak in OHCs^7,13,14^, suggesting that *Myo7a-N* is driven by an alternative promoter. A previous study in zebrafish identified two regulatory ‘blocks’ upstream of the *Myo7a* gene that drive hair cell–specific expression of reporter genes^12^. Our recent findings suggest that one of these blocks corresponds to the first exon of a previously uncharacterized *Myo7a* isoform we named *Myo7a-N*^15^, while the other is located just upstream, consistent with the location of a putative *Myo7a-N* promoter. In the future, it will be important to investigate how *EnhancerA* interacts with these isoform-specific promoters to regulate MYO7A expression in a cell type- and isoform-specific manner.

### MYO7A tunes MET gating

It has been proposed that MYO7A acts as a motor protein that provides tension to the MET complex at UTLDs^34–36^. However, due to the disruption of hair bundle development in MYO7A depletion models, it has been challenging to isolate its specific role in MET function. Previously, we employed the *Myo7a-*Δ*C* model to examine the effects of severe MYO7A depletion specifically in IHCs^7^. In this model, reduction of MYO7A at UTLDs led to decreased resting P_o_ and slowed onset of MET currents, supporting the hypothesis that MYO7A contributes to tensioning the MET complex.

In the current study, the Δ*EnhancerA* model serves as a tool to test MYO7A function, as it reduces expression of all MYO7A isoforms in both IHCs and OHCs. Notably, the tonotopic gradient of MYO7A depletion in this model enables investigation of MET function across a range of MYO7A expression levels. At P4-5, MYO7A reduction is most severe in basal OHCs, where we observed a significant decrease in MET resting P_o_, while peak current amplitude and adaptation kinetics remained unaffected. In contrast, apical OHCs, which exhibited only mild MYO7A depletion, showed no measurable changes in MET properties. Together, these results reinforce our previous findings that MYO7A depletion leads to a reduction in tip-link tension^7^.

Notably, a recent study reported that MYO7A loss in mature hair cells (P20–P30) led to reduced bundle stiffness and diminished peak MET currents, without affecting resting Po^5^. The contrast with our findings at P4-5 may be due to developmental stage-specific functions of MYO7A. During early postnatal stages, MYO7A may play a more active role in generating tip-link tension and supporting MET complex assembly, whereas in mature cells, its function may shift toward structural stabilization. Collectively, these results underscore the dynamic, isoform- and stage-dependent contributions of MYO7A to hair cell MET that warrants further research to gain a more complete picture.

### Other regulatory mechanisms that drive *Myo7a* expression

We propose that *EnhancerA* functions as a shared scaffold for transcriptional activation in both IHCs and OHCs, with isoform specificity determined by the local presence or absence of trans-acting factors. Using publicly available ChIP-seq datasets, we identified a SIX2 binding site within *EnhancerA*^24^. Our qPCR and immunofluorescence analyses demonstrate that conditional deletion of SIX2 leads to a reduction in MYO7A expression, with the MYO7A-N isoform being more strongly affected than other isoforms. These results suggest a potential regulatory role for SIX2 in isoform-specific control of MYO7A expression: in OHCs, particularly in basal regions where SIX2 expression is highest, SIX2 may cooperate with *EnhancerA* to drive *Myo7a-N* expression via a potential *Myo7a-N* promoter, thereby establishing an increase in *Myo7a-N* levels toward the cochlear base. In contrast, in IHCs, which express little or no SIX2, *EnhancerA* activity may preferentially be coupled with the canonical promoter of *Myo7a-C* via other transcription factors. This combinatorial mechanism, where a broadly active enhancer integrates input from spatially restricted transcription factors, offers a potential explanation for the observed isoform gradients.

While our data implicate SIX2 in regulating MYO7A expression, its precise contribution remains unclear, as we did not test direct binding of SIX2 to *EnhancerA*. Meanwhile, the residual MYO7A expression observed in Δ*EnhancerA* mice suggests the existence of additional regulatory mechanisms governing *Myo7a* expression. Notably, a recent parallel study independently identified the *EnhancerA* region (referred to as E3 in their study), along with several additional *cis*-regulatory elements within the *Myo7a* locus^37^. In their study, E3 drives reporter gene expression only in combination with other *Myo7a* enhancers, suggesting that *Myo7a* expression is regulated by a more complex cooperative *cis/trans*-regulatory element network.

### SIX2 is dispensable for hair cell development but required for maintenance

SIX family transcription factors are known to be critical for cell proliferation and differentiation, which is important for kidney development^27–29,38^. SIX1 and SIX2 share high similarity in amino-acid sequence, structural domains and DNA-binding motif. Both have been implicated in auditory system development, with SIX1 being extensively characterized for its essential role in early inner ear development^39–43^. A recent human genetic study associates SIX2 haploinsufficiency with mid-high frequency hearing loss^31^, however, the molecular function of SIX2 in the cochlea has been less defined.

In this study, we demonstrated that SIX2 is not required for hair cell development. However, SIX2 contributes to the maintenance of mature hair cells, as hair cell-specific deletion using the *Myo15-cre* mouse leads to progressive loss of OHCs beginning in the basal region of the cochlea by 10 weeks of age, accompanied by hair bundle degeneration and high-frequency hearing loss. Deletion of SIX2 reduces overall Myo7a expression, with the Myo7a-N isoform showing a more pronounced reduction, while Myo7a-C exhibits only a modest, statistically insignificant decrease. Notably, it is an intriguing observation that postnatal deletion of SIX2 results in a relatively mild phenotype, despite the strong enrichment of SIX2 expression in OHCs, suggesting redundancy and compensation by other transcription factors. As a future direction, it will be important to identify downstream targets of SIX2 using RNA sequencing. In addition, because SIX2 is also expressed in the GER and other supporting cell types, it will be informative to delete SIX2 using an inner ear sensory epithelium-specific Cre driver and perform single-cell RNA sequencing to characterize its downstream targets in non-hair cell populations.

As mentioned above, SIX2 likely operates within a wider transcriptional network involving other regulators such as ATOH1^44^, POU4F3^45^, and GFI1^46^. While other transcription factors initiate hair cell differentiation, SIX2 may refine OHC identity by regulating genes that are required for hair cell maturation and maintenance. Meanwhile, EYA1/2 are known transcription co-factors that interact with SIX2 for gene expression, and EYA2 showed tonotopic hair-cell specific expression^47^. Interactions among these transcription factors are likely essential for shaping MYO7A isoform diversity and for establishing the tonotopic architecture of cochlear hair cells.

### Implications for isoform-specific gene therapy

Our findings have potential translational implications, given the compact size of *EnhancerA* and its unique spatiotemporal regulatory pattern. Meanwhile, *EnhancerA* and SIX2-responsive elements could be leveraged to design synthetic promoters that enable targeted transgene expression in hair cells, or even regionally along the cochlear axis. This spatiotemporal control would be particularly valuable for treating late-onset, genetic forms of deafness, in which precise regulation of expression timing and location is critical for therapeutic efficacy.

In summary, this study identifies *EnhancerA* and SIX2 as regulators of MYO7A isoform expression in the cochlea, linking transcriptional control to regional and cell type-specific functional specialization. These findings deepen our understanding of cochlear gene regulation and lay the groundwork for targeted, isoform-aware therapies for sensorineural hearing loss.

**Supplementary Figure 1:**
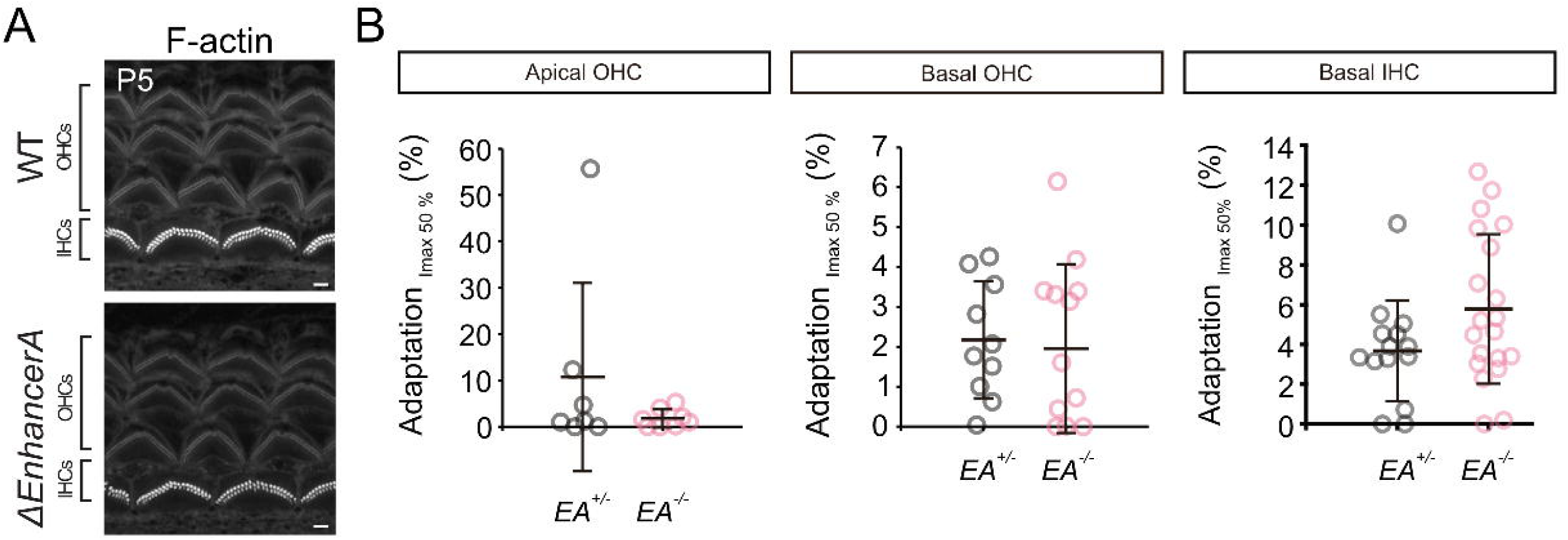
A. Immunofluorescence images of WT and Δ*EnhancerA* mice at P5 in the basal region of the cochlea. Hair bundle morphology in Δ*EnhancerA* mice is comparable to that of WT mice. (Scale bar: 1 µm). **B**. Summary plots of the adaptation for apical OHCs, basal OHCs and basal IHCs. Slow adaptation was measured at the 50% point of the total peak current. Whiskers of the plot represent for Mean ± SD. Number of cells (animals): Apical *EA^+/+^* = 8 (8), Basal *EA^+/+^* 13 (13), Apical *EA^−/−^* 7 (7), Basal *EA^−/−^* 10(10).

## 7. Methods

### Animal care and handling

The care and use of animals for all experiments described conformed to NIH guidelines. Experimental mice were housed in a 12:12□hour light:dark cycle with free access to chow and water in standard laboratory cages in a temperature and humidity-controlled vivarium. The protocols for the care and use of animals were approved by the Institutional Animal Care and Use Committees at the University of Virginia, the University of Colorado Denver, and the National Institute on Deafness and Other Communication Disorders. All the institutions mentioned above are accredited by the American Association for the Accreditation of Laboratory Animal Care. C57BL/6J (Bl6, from Jackson Laboratory, ME, USA) mice and sibling mice served as control mice for this study. Neonatal mouse pups (P0–P5) were sacrificed by rapid decapitation. Mature mice were euthanized by CO_2_ asphyxiation followed by cervical dislocation.

### Generation of Mouse strains for this study

For CRISPR/Cas mediated generation of the mouse models, we used the online tool CRISPR (http://crispor.tefor.net/crispor.py) to select suitable target sequences to ablate *Myo7a* isoforms. Single-guide (sg)RNAs are purchased from IDT (Integrated DNA Technologies, IA, USA). To produce genetically engineered mice, fertilized eggs were coinjected with Cas9 protein (PNA Bio, 50□ng/µl) and the sgRNA (30□ng/µl). Two-cell stage embryos were implanted on the following day into the oviducts of pseudopregnant ICR female mice (Envigo). Genotyping was performed by PCR amplification of the region of interest. Target sequences, sgRNAs, repair templates, and genotyping primers are listed below:

#### ΔEnhancerA mice

To ensure the large deletion of *EnhancerA*, a pair of guide RNA surrounding the *EnhancerA* was injected. sgRNA1: 5’-ACAACAGCGACUUCCCUGCAUGG-3’, sgRNA2: 5’-CCTACGTTCCTAG CCAGTCTCGC-3’. Genotyping primers: Forward: 5’-ACCATGACCCCATAAGGTTC-3’, W T reverse: 5’-GGGATGAATCATCACTCCTGC-3’, Mutant Reverse: 5’-AGTGTTCTCCCGA GAGCCTTTC-3’.

#### Six2-HA mouse

sgRNA: 5’-AGAGCTCTGTTCGCCTCCGG-3’. Repair template: GCGGCGCGGACCC ACTGCAGCATCACCACAGCCTGCAGGACTCCATACTCAACCCCATGTCGGCCAACCTTGTGGACCTGGGCTCCTACCCATACGATGTTCCAGATTACGCTTAGAGCTCTGTTCGCCTCCGGGCGCTTGCTTTCCGTGCTTGATGAAGCGGTGTGGGAGGGCGATAGATTCTGGAGTCTGTGCCGCGT. Genotyping primers: Forward: 5’-TCCATACTCAACCCCATGTCGG-3’, Reverse: 5’-CCAAAGGATACCGAGCAGACCA-3’.

#### Gfi1-cre

*Gfi1^tm^*^1^*^(cre)Gan^*(*Gfi1-cre*) mouse line is Dr. Lin Gan at the University of Rochester. A targeting construct was used to replace the coding sequence of *Gfi1* from the translation initiation codon through exon 5 with the coding sequence of cre recombinase. Genotyping primers: CRE-Forward: 5’-GGGATAACGGACCAGTTG-3’, Reverse: 5’-GCCCAAATGTTG CTGGATAGT-3’. *Gfi1-cre* WT allele genotyping primer: Forward: 5’-GGGATAACGGACCAGTTG-3’, Reverse: 5’-CCGAGGGGCGTTAGGATA-3’ (More information: https://www.informatics.jax.org/allele/MGI:4430258#:∼:text=Mutation%20details%3A%20A% 20targeting%20construct,coding%20sequence%20of%20cre%20recombinase.)

#### Myo15a-Cre

To generate a *Myo15a-Cre* knockin (KI) mouse line, CRISPR/Cas9-mediated genome engineering was used to insert a “Cre-rBG (rat beta-globin intron) polyA” cassette upstream of the translational start codon (ATG) in exon 2 of the mouse *Myo15* gene (NCBI Reference Sequence: NM_010862.2), which is located on chromosome 11 and contains 66 exons (Ensembl Transcript: ENSMUST00000071880). The cassette was placed under the control of the endogenous *Myo15a* promoter to ensure physiologically relevant Cre expression. Homology arms (∼2–3 kb) flanking the insertion site were amplified by high-fidelity PCR using BAC clone RP23-329M3 as a template and assembled with the Cre-rBG cassette into the targeting vector. For genome editing, Cas9 protein, guide RNAs (gRNAs), and the targeting vector were co-injected into fertilized C57B L/6J one-cell embryos. Two gRNAs were designed to flank the insertion site with mini mal predicted off-target activity: gRNA-A1 (GAGGGCCACCATGGCGGATG) matching th e forward strand and gRNA-B1 (TTTCTTCTCCTCATCCGCCA) matching the reverse strand. Injected embryos were transferred into pseudopregnant females, and resulting pups were genotyped by PCR using genomic DNA from tail biopsies. Correct integration of the cassette was confirmed by Sanger sequencing across the homology arm–cassette junctions. The mouse line was generated by Cyagen Biosciences (Santa Clara, CA), and correctly targeted founders were bred with C57BL/6J animals to establish a stable *Myo15a-Cre* line.

Genotyping primers: CRE-Forward: 5’-GGGATAACGGACCAGTTG-3’, Reverse: 5’-GCCCAAATGTTGCTGGATAGT-3’. *Myo15a-Cre* WT allele genotyping primer: Forward: 5’-GGTCTCCAAGCAGTGTCTCCAA-3’, Reverse: 5’- CTTGAGGCTCCGTTTGGGTTTC -3’ *Six2::cre-egfp* mice is purchased from The Jackson Laboratory (Strain #:009606), and *Six2^flox/flox^* are generated by previous studies from other labs. The genotyping methods are provided by their corresponding reference^48,49^.

### Quantitative polymerase chain reaction (qPCR)

qPCR primers were designed to detect either *pan-Myo7a* transcripts or *Myo7a* isoform-specific transcripts to evaluate their expression levels. Total RNA was extracted using TRIzol reagent (15596026, ThermoFisher Scientific). Reverse transcription was performed using the SuperScript™ IV Reverse Transcription system (18090010, ThermoFisher Scientific). qPCR reactions were conducted using iTaq Universal SYBR Green Supermix (1725121, Bio-Rad, CA, USA) on a CFX Opus 384 Real-Time PCR System (12011319, Bio-Rad) for fluorescence measurement. Following analysis and plotting is done by R (version 4.4.1).

Primers: *pan-Myo7a*: forward: 5’-CCTTCTCATTCGCTACCGGG-3’, reverse: 5’-CCGG TTGTTGCGTTTCATGT-3’; *Myo7a-C*: forward: 5’-CCGGTTGTTGCGTTTCATGT-3’, reverse: 5’-CCGACTCCCCGCTGATAATAC-3’ ; *Myo7a-N*: forward: 5’-GTGCGAAACTTGAACCA AACG-3’, reverse: 5’-CCGACTCCCCGCTGATAATAC-3’ ; *Myo7a-S*: forward: 5’-ACAACA GCGACUUCCCUGCAUGG-3’, reverse: 5’-CCGACTCCCCGCTGATAATAC-3’ ; *Atp8b1*: forward: 5’-ACGGCCGGTGGTCTTACATA-3’, reverse: 5’-GCTTGTCACTCACGTCCTGAT-3’ ; *Tmc1*: forward: 5’-GTGGGCCCTTCAGTGGTAAA-3’, reverse: 5’-TTTCTGGCCCTTGG CAGTAG-3’ ; *Tmc2*: forward: 5’-GCCGGAAGGATCCTGCTAAG-3’, reverse: 5’-CAAGGA AGTGGACCTGGGTT-3’.

### Immunofluorescence

Inner ear organs were fixed in 3% paraformaldehyde (PFA, Electron Microscopy Sciences, PA) immediately after dissection for 20□min. Samples were washed three times with phosphate-buffered saline (GIBCO^®^ PBS, Thermo Fisher Scientific, Waltham, MA) for 5□min each. After blocking for 2□h with blocking buffer (1% bovine serum albumin, 3% normal donkey serum, and 0.2% saponin in PBS), tissues were incubated in blocking buffer containing primary antibody at 4□°C overnight. The following antibodies were used in this study: rabbit polyclonal Myosin-VIIa antibody (catalog#: 25-6790, Proteus Biosciences Inc, Ramona, CA. 1:100), mouse monoclonal Myosin-VI antibody (A-9, Santa Cruz, 1:100), rabbit anti-HA antibody (C29F4, Cell Signaling Technologies, Catalog #: #3724, Danvers, MA). Fluorescence imaging was performed using a Leica Stellaris 5 Confocal Microscope Platform or Zeiss LSM880 with AiryScan.

### Hearing tests in mice

Mice were anesthetized with a single intraperitoneal injection of 100□mg/kg ketamine hydrochloride (Fort Dodge Animal Health) and 10□mg/kg xylazine hydrochloride (Lloyd Laboratories). ABR and DPOAE were performed in a sound-attenuating cubicle (Med-Associates, product number: ENV-022MD-WF), and mice were kept on a Deltaphase isothermal heating pad (Braintree Scientific) to maintain body temperature.

ABRs of Δ*EnhancerA* mice and their BL6 controls were conducted on the equipment purchased from Intelligent Hearing Systems (Miami, Fl). Recordings were captured by subdermal needle electrodes (FE-7; Grass Technologies). The non-inverting electrode was placed at the vertex of the midline, the inverting electrode over the mastoid of the right ear, and the ground electrode on the upper thigh. Stimulus tones (pure tones) were presented at a rate of 21.1/s through a high-frequency transducer (Intelligent Hearing Systems). Responses were filtered at 300–3000□Hz, and threshold levels were determined from 1024 stimulus presentations at 8, 11.3, 16, 22.4, and 32□kHz. Stimulus intensity was decreased in 5–10□dB steps until a response waveform could no longer be identified. Stimulus intensity was then increased in 5□dB steps until a waveform could again be identified. If a waveform could not be identified at the maximum output of the transducer, a value of 5□dB was added to the maximum output as the threshold.

DPOAEs of the same group of WT and *Myo7a-*Δ*C* mice were recorded. While under anesthesia for ABR testing, DPOAE was recorded using SmartOAE ver. 5.20 (Intelligent Hearing Systems). A range of pure tones from 8 to 32□kHz (16 sweeps) was used to obtain the DPOAE for the right ear. DPOAE recordings were made for *f*2 frequencies from 8.8 to 35.3□kHz using a paradigm set as follows: *L*1□=□65□dB, *L*2□=□55□dB SPL, and f_2_/f_1_□=□1.22.

ABRs of *Myo15a::cre; Six2^fl/fl^* mice and its corresponding control are conducted on RZ6 Multi-I/O Processor (Tucker Davis, FL 32615 USA) with Medusa4Z ABR Amplifier. Responses are averaged by 512 stimulus sweeps presentations at 5, 10, 16, 24, 32, 40 and 48□kHz. Same ABR threshold determination strategies mentioned above are applied.

### Whole-cell voltage-clamp electrophysiology recordings

#### Electrophysiological recordings

P4-P5 Δ*EnhancerA* homozygous and heterozygous mice were used for whole-cell patch clamp experiments. Recordings were collected on the first or second row of outer hair cells using an Axon 200B amplifier or a Multiclamp 700B (Molecular Devices). Initially, a patch pipette filled with extracellular solution was used to clean the epithelium, and 2-3 adjacent cells near the target cell were removed. Subsequently, a patch pipette filled with intracellular solution was used to perform the whole-cell patch clamp on the cell. Apical perfusion was continuously maintained until a gigaseal was achieved. After breakthrough of the patch membrane to enter whole-cell configuration, a minimum waiting time of 5 minutes was allowed for the intracellular solution to equilibrate. A fluid jet, using thin-walled borosilicate pipettes (World Precision Instrument), was then applied to deliver step-like force stimuli to the hair bundle. Experiments were performed at 18-22°C. Whole-cell currents were filtered at 10 kHz and sampled at 50 kHz using USB-6356 or USB6366 (National Instruments) controlled by jClamp (SciSoft Company). All experiments used −80 mV holding potential unless otherwise noted and did not account for the liquid junction potential.

#### Hair-bundle stimulation and motion recording

For the electrophysiological recordings, hair bundles were stimulated with a custom three-dimensional—printed fluid jet driven by a piezoelectric disc bender (27 mm, 4.6 kHz; Murata Electronics or PUI Audio). Thin-wall borosilicate pipettes were pulled to tip diameters of 5 to 15 µm, filled with extracellular solution, and mounted in the fluid-jet stimulator. The piezoelectric disc bender was driven by waveforms generated using jClamp. Stimulus waveforms were filtered using an eight-pole Bessel filter at 1 kHz (L8L 90PF, Frequency Devices Inc.) and variably attenuated (PA5, Tucker Davis) before being sent to a high-voltage/high-current Crawford amplifier to drive the piezoelectric disc bender. During stimulations, videos were taken of the hair-bundle motion with high-speed imaging at 10,000 frames per second using a Phantom Miro 320s or Veo 410L camera (Vision Research) when illuminated with a TLED+ centered at 530 nm (Sutter Instruments). Videos were saved for each stimulation and analyzed offline. Hair bundle displacement was determined as described previously^50–52^. Briefly, hair bundle position was extracted using a Gaussian fit to a high-pass–filtered hair-bundle image for a given vertical row of pixels in the image.

#### Analysis

All data were analyzed offline using jClamp (SciSoft), Matlab (MathWorks), Excel (Microsoft), and Prism 7 (GraphPad). Figures were generated using Matlab, Prism 7, and Adobe Illustrator.

Electrophysiological data analysis: The resting open probability of MET channels (defined as P_open_, or P_o_) was determined from sine wave stimulation of the hair bundle and calculated using Eq. (2), where I_max_ is the current elicited during maximum positive stimulation, I_leak_ is the current remaining during maximum negative stimulation, and I_resting_ is the resting mechanosensitive current (defined as the current in the absence of stimulation, subtracted by I_leak_). We assumed we could observe a P_o_ of 0% and 100% during the maximum negative and positive stimulations, respectively,

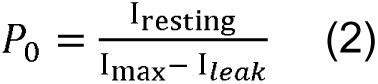

For mechanical stimulus steps, adaptation time constant fits were obtained at ∼50% peak current using a double exponential equation

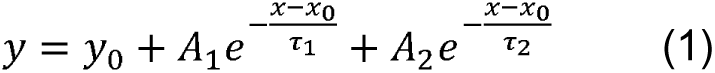

where τ_1_ and τ_2_ are the decay constants, and A_1_ and A_2_ are the respective amplitudes; the hair-bundle displacement was also fit with the same equation starting at a time after the force stimulus plateaued, which was 0.5 ms after stimulus onset. All fits were automated in MATLAB.

### Scanning electron microscopy

Adult mice were euthanized by CO2 asphyxiation before intracardiac perfusion with 2.5% glutaraldehyde (Electron Microscopy Sciences, Hatfield, PA) and 2% PFA. The otic capsule was dissected and incubated in postfixation buffer at 4□°C overnight (2.5% glutaraldehyde, in 0.1□M cacodylate buffer, with 3□mM CaCl2). For neonatal mouse pups, the samples were dissected and treated with postfixation buffer immediately. The otic capsules from adult mice were incubated for two weeks in 4.13% EDTA for decalcification and then further dissected to expose the organ of Corti. Samples underwent the OTOTO procedure and were dehydrated using gradient ethanol and critical point drying. After sputter coating with platinum, the samples were imaged on Zeiss Sigma VP HD field emission SEM using the in-lens secondary electron detector.

## Acknowledgement

We thank Wenhao Xu and Daniel Grigsby at Genetically Engineered Murine Model Core (University of Virginia, RRID:SCR_025473) for generating mouse models used in this study. We thank Sijie Hao from Advanced Microscopy Facility (University of Virginia, RRID: SCR_018736) for the support of SEM imaging. We thank Ruliang Jiang (Cincinnati Children’s Hospital Medical Center) for providing *Six2^flox/flox^* mice. We thank Maria Luisa S. Sequeira Lopez (University of Virginia) for providing the *Six2::cre-egfp* mice.

Funding source: J.B.S was supported by NIH grant RO1DC018842. Zeiss SEM of Advanced Microscopy Facility was supported by NIH SIG grant 1S10OD011966-01A1. The Titan Krios of Molecular Electron Microscopy Core was purchased with NIH grant SIG S10-RR025067. Y.R.K was supported by the National Research Foundation of Korea (NRF) grant funded by the Korea government (MSIT) (RS-2025-24535196).

## Author contributions

Conceptualization: S.L., A. W.P., J.B. Shin; Methodology: S.L., S.J., Y.R.K., C.K., A.W.P., J.B. Shin; Experiments: S.L., S.J., Y.R.K., A.G., F.L.; Writing—review & editing: S.L., S.J., A. W.P., J.B. Shin; Supervision: U.K., A. W.P., J.B. Shin.

## Notes

### Competing Interest Statement

The authors have declared no competing interest.

### Summary of Updates

Figure 3-6 revised. Updated results for Gfi1-cre and Myo15a-cre induced Six2 conditional knock-out mice.

